# Quantification of *Brucella abortus* population structure in a natural host

**DOI:** 10.1101/2020.11.12.380766

**Authors:** Aretha Fiebig, Catherine E. Vrentas, Thien Le, Marianne Huebner, Paola M. Boggiatto, Steven C. Olsen, Sean Crosson

**Affiliations:** Department of Microbiology and Molecular Genetics, Michigan State University, East Lansing, MI, 48824, USA; Infectious Bacterial Diseases Research Unit, National Animal Disease Center, Agricultural Research Service, U.S. Department of Agriculture, Ames, IA, 50010, USA; Department of Statistics and Probability, Michigan State University, East Lansing, MI, 48824, USA

## Abstract

Cattle are natural hosts of the intracellular pathogen, *Brucella abortus*, which inflicts a significant burden on the health and reproduction of these important livestock. The primary routes of infection in field settings have been described, but it is not known how the bovine host shapes the structure of *B. abortus* populations during infection. We utilized a library of approximately 10^6^ uniquely barcoded *B. abortus* strains to temporally and spatially quantify population structure at the strain level during colonization of cattle through a natural route of infection. Introducing 10^8^ bacteria from this barcoded library to the conjunctival mucosa resulted in expected levels of local lymph node colonization at a one-week timepoint. We leveraged variance in strain abundance in the library to demonstrate that only 1 in 10,000 brucellae introduced at the site of infection reached the parotid lymph nodes. Thus, cattle restrict the overwhelming majority of *B. abortus* introduced via the ocular conjunctiva at this dose. Individual strains were spatially restricted within the host tissue, and the total *B. abortus* census was dominated by a small number of distinct strains in each lymph node. These results define a bottleneck that *B. abortus* must traverse to colonize local lymph nodes from the conjunctival mucosa. The data further support a model in which a small number of spatially isolated granulomas founded by unique strains are present one-week post infection. These experiments demonstrate the power of barcoded transposon tools to quantify infection bottlenecks and to define pathogen population structure in host tissues.

**Significance statement:** Understanding microbial population dynamics during infection has important implications for disease management, transmission and pathogen evolution. A quantitative analysis of microbial population structure requires the ability to track individual strains. We used a pool of individually barcoded strains to measure changes in *Brucella abortus* population structure during infection of bovine hosts via the ocular conjunctiva, a natural route of entry. Cattle exert a severe bottleneck on the bacterial population entering through the conjunctival mucosa such that individual cells have a 0.0001 probability of colonizing a local draining lymph node. The populations in lymph nodes, even on different sides of the same animal, are distinct and dominated by a small number of highly abundant, spatially distinct clones.

## Introduction

*Brucella abortus* commonly infects domesticated cattle (2) and is an etiologic agent of brucellosis, a global zoonotic disease that inflicts a significant burden on human and animal health and agricultural production (3). Our understanding of *Brucella* infection biology and host response has been greatly informed by *in vitro* studies of infection in tissue culture, or by *in vivo* modeling of brucellosis in rodents. Mouse models are particularly well developed and have proven useful in defining features of mammalian immune protection and in testing new vaccines and brucellosis therapies (4). However, unlike cattle, mice are not natural *Brucella* hosts. Moreover, the experimental routes of infection employed in laboratory studies of the mouse do not typically reflect natural routes of host entry that occur in *bona fide* mammalian hosts under field conditions. New studies of *B. abortus* using natural hosts infected through natural routes are therefore crucial as we work to advance understanding of the infection biology of this important group of pathogens.

Bacteriological analyses of infected cattle provide evidence that *B. abortus* entry through the ocular conjunctiva and upper respiratory passage is common under field conditions (5). Entry also occurs via mucosal tissue of the oral cavity, the respiratory tract, or the gastrointestinal tract (6–9). While invasion of the conjunctiva and other mucosal tissue induces an inflammatory response in the submucosa of cows (10), it is clear that some fraction of *B. abortus a)* survive this inflammation after traversing the host barrier, *b)* enter the lymphatic system, and *c)* begin to colonize the parotid lymph node within one week of infection (1, 5, 10, 11). The fraction of infecting *B. abortus* that eventually colonize lymph nodes though these natural routes of entry remain largely undefined.

Population bottlenecks are common during infection through natural routes as host barriers and immune functions serve to minimize pathogen entry and survival (12). Studies involving pools of strains with a limited number of unique tags have been utilized to assess bottlenecks during infection in model systems (eg. (13–16)). Understanding the magnitude of bottlenecks is important in developing models of pathogen evolution and microbial adaptation to the host (17, 18). Knowledge of bottlenecks also greatly informs the design and interpretation of transposon-insertion sequencing (Tn-seq) studies aimed at discovering genes required for survival in the animal host (19). In this study, we aimed to define the magnitude of the bottleneck during *B. abortus* infection of its natural bovine host via a natural mucosal route of infection, and to quantify *B. abortus* population structure in the host lymphatic tissue after infection.

We previously generated a highly diverse pool of approximately 10^6^ *B. abortus* transposon mutant strains (20, 21). A 20-base pair barcode present in each transposon mutant (22) provides a unique identification tag for each *B. abortus* strain and thus enables strain-level quantification of *B. abortus* populations. In this study, we used this diverse pool of individually tagged strains to study changes in *B. abortus* population structure upon infection through the conjunctival mucosa in cattle. Our study characterizes infection bottlenecks along the conjunctival route of entry, defines spatial distributions of clonally-derived *B. abortus* strains in lymph nodes that drain the ocular and oral cavities, and demonstrates the feasibility and power of Tn-seq approaches to study *B. abortus* populations in a bovine infection model.

## Results

### Reproducible infection of calves via the conjunctival route with a *Brucella* abortus Tn-*Himar* mutant library

Three-to four-month-old calves are reproducibly infected after experimental conjunctival exposure to 5 x 10^8^ CFU of *B. abortus* (1). The ocular conjunctiva is reported to be a natural route of host entry in the field (5), and this method of infection is often used for vaccine challenge studies. Thus, we administered 10^8^ CFU from our barcoded *B. abortus* Tn-*himar* mutant library to the right and left ocular conjunctiva of six calves between 5 and 9 months of age (**Table 1**). Infected animals were necropsied 8-or 9-days post-infection to assess *B. abortus* mutant populations in the parotid and retropharyngeal lymph nodes (**Fig 1A**). Homogenates of the right and left parotid lymph nodes of each of the 6 calves yielded approximately 10^5^ CFU/g of tissue (**Fig 1B**). We further assessed colonization of the left retropharyngeal lymph nodes, which are distally draining compared to the parotid, and noted high variability in the extent of colonization between animals. Populations of *B. abortus* were isolated from retropharyngeal homogenates of only 3 of the 6 infected animals at our 8-to 9-day time point. This result is consistent with the work of Meador and colleagues (1), which demonstrated that *B. abortus* introduced via the conjunctival route is detected in the retropharyngeal lymph node between 7 and 21 days post-infection.

**Fig 1.**
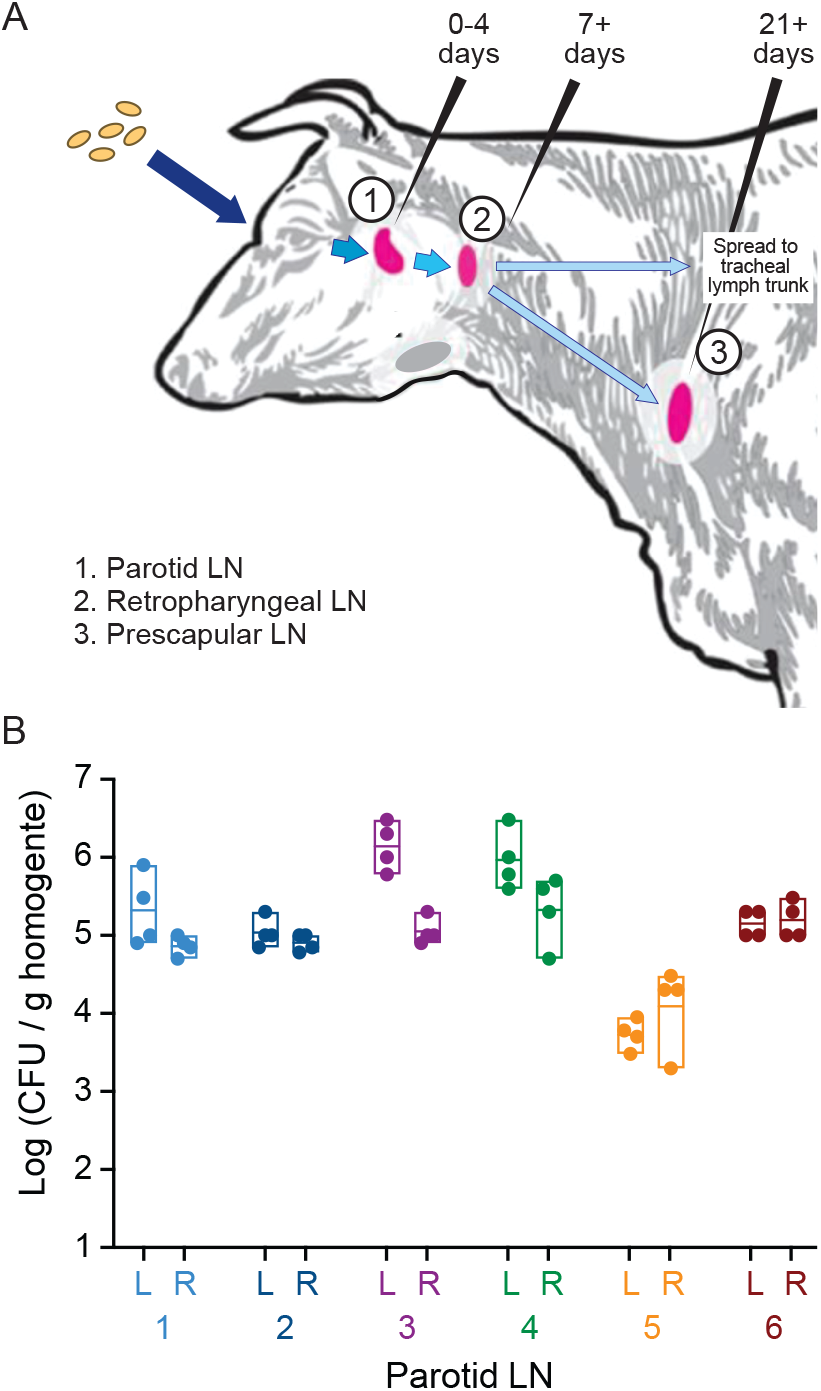
Colonization of bovine parotid lymph nodes by the barcoded Tn-*himar Brucella abortus* strains. **A.** A barcoded pool of *B. abortus Tn-himar* strains (tan ovals) were inoculated into the left and right ocular conjunctiva of six calves. These bacterial cells disseminate to the lymph nodes (LN) of the head and neck (pink). *B. abortus* can be detected in (1) the parotid LN at 0-4 days post-infection, (2) the retropharyngeal LN at 7-21 days post-infection, and (3) the prescapular LN at timepoints at least 21 days after infection (1). **B.** Expected levels of colonization were observed in the parotid LN 8-9 days post infection. Parotid LN from the left (L) and right (R) sides of the infected animals were removed, quartered and homogenized. Dots represent colony forming units (CFU) per gram of tissue that were enumerated from each homogenized lymph node quadrant. Boxes indicate the minimum, maximum and mean titer of the quadrants in each parotid LN of each animal (numbered 1-6).

**Table 1.** Calves utilized in the study.

| Animal | Sex | Age at time of infection (months) | Days post-infection at necropsy |
| --- | --- | --- | --- |
| 1 | Bull | 5 | 8 |
| 2 | Bull | 5 | 9 |
| 3 | Heifer | 9 | 9 |
| 4 | Heifer | 9 | 8 |
| 5 | Heifer | 5 | 8 |
| 6 | Heifer | 5 | 9 |

### Amplification and sequencing of barcodes from *B. abortus* strains isolated from bovine lymph nodes

To assess the *B. abortus* Tn-strain distribution in host tissue, lymph node homogenates were spread on TSA-SK agar, the same medium used to outgrow the library for inoculation. Transposon barcodes were PCR amplified from genomic DNA extracted from these plated samples, and barcode amplicons were sequenced and counted. The numbers of sequence reads and unique barcodes detected in each sample are detailed in **Dataset S1**. Using this approach, barcodes were reliably amplified, identified, and tallied across samples. Each lymph node fragment and inoculum sample yielded a similar number of total reads (2 - 5 x 10^6^), indicating similar sampling depths for all tissue samples (**Fig 2A**).

**Fig 2.**
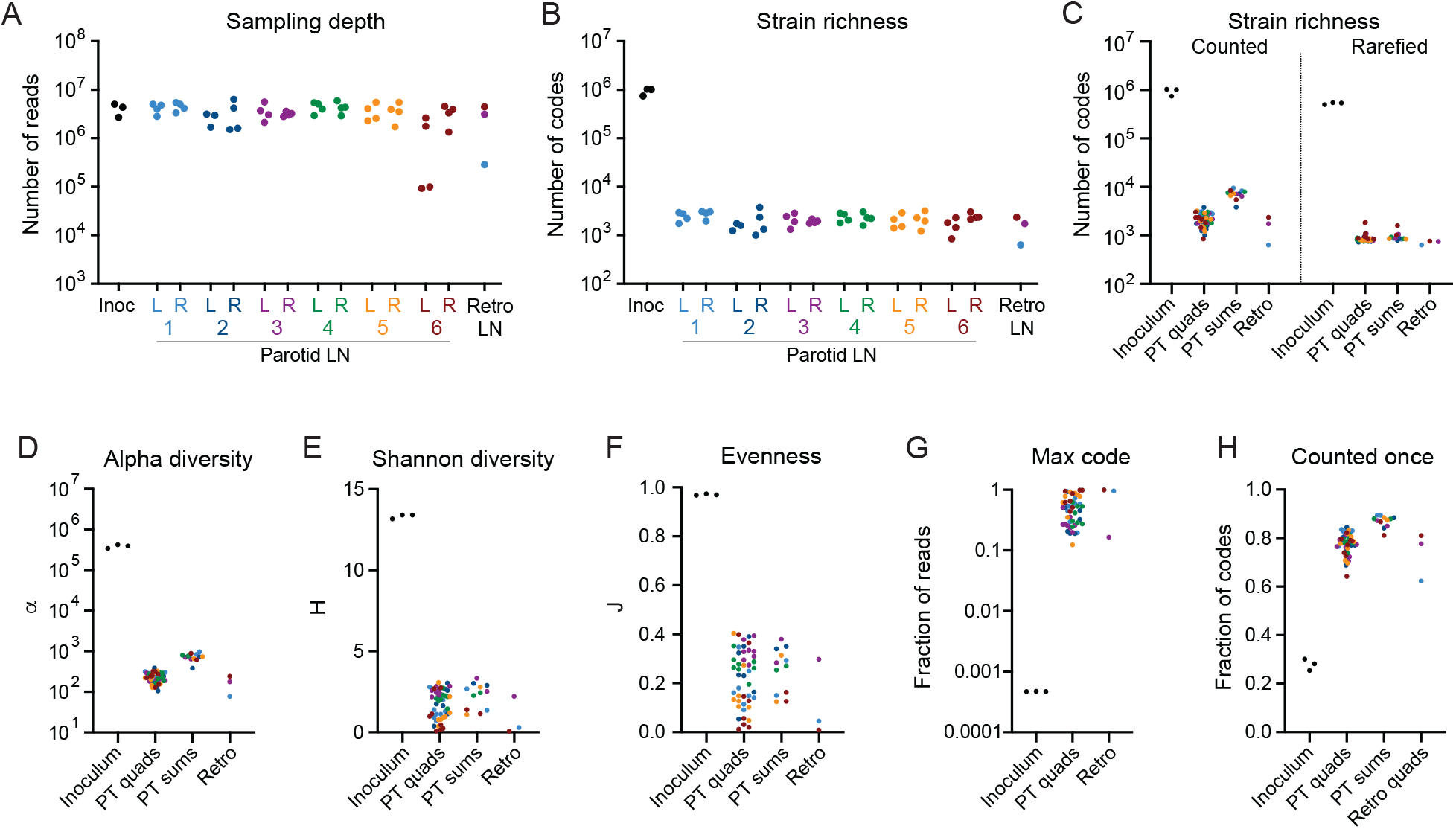
Loss of strain diversity in lymph node samples. **A.** Number of barcode sequencing reads from each sample. Each dot represents a sample from the inoculum (inoc), a parotid lymph node (LN) quadrant from the left (L) or right (R) side of each animal (numbered 1-6), or a colonized retropharyngeal lymph node (Retro LN). **B.** Number of unique barcodes (i.e. strains) counted in each sample. Each dot represents a sample as in A. **C.** Number of codes counted in each type of sample before (left) and after (right) rarefaction to normalize for differences in sequencing depth. **D, E, F.** Fisher’s alpha diversity, a, Shannon diversity, H, and Shannon evenness, J, calculated for each sample. **G.** Fraction of reads represented by the most abundant barcode in each sample. **H.** Fraction of barcodes counted only once in each sample. PT quads: individual quadrants from parotid LNs. PT sums: the sum of barcode counts for the quadrants of each parotid LN. Retro; fragment from colonized retropharyngeal LN. Points are colored by animal as in **Fig 1**.

### The Tn-Himar mutant pool is a diverse and evenly distributed population of B. abortus strains

The distribution of barcodes in the *B. abortus* mutant pool (i.e. the inoculum) reflects a diverse and evenly distributed set of unique strains. We identified 0.7 – 1.0 x 10^6^ barcodes in each of three inoculum aliquots, which collectively represent 1.5 x 10^6^ unique codes (**Fig S5**). We previously mapped 5 x 10^5^ barcodes to distinct positions in the *B. abortus* genome with high confidence (21), and in this present study, we captured most of the previously mapped barcodes in the inoculum (**Fig S5**). Within each inoculum sample, barcode abundances were evenly distributed with a small tail of more abundant codes (**Fig S6**). These samples contained few codes represented by 100 or more reads and, accordingly, our filtering approach outlined in Materials and Methods and **Fig S2** removed only 0.2 % of reads and 0.7% of the codes from these samples (**Fig S3**, **Dataset S1**). The apparent strain diversity of the inoculum samples may therefore be modestly inflated by sequencing errors, which are expected in approximately 1% of the barcode reads (23–25) (and examples in **Fig S1**).

### *B. abortus* strain diversity in lymph nodes is highly reduced relative to the inoculum

We next used our barcode sequence data sets to assess changes in *B. abortus* population structure when *B. abortus* enters the host via the conjunctival route. Though we administered 10^8^ CFU of a library containing at least 10^6^ unique strains to the right and left ocular conjunctiva of each calf, less than 10^4^ unique barcodes were recovered from each intact lymph node on the right and left sides of the animal (**Fig 2C**). In each lymph node quadrant we detected an average of 2 × 10^3^ unique barcodes (**Fig 2B-C**). To determine if differences in apparent strain diversity between host tissue samples were due to variability in sequencing depth (**Fig S3**), we randomly selected 10^6^ reads from each sample for barcode analysis. This rarefaction to normalize for differences in sampling depth indicates a nearly uniform number of strains in each lymph node (**Fig 2C**). We conclude that the number of unique strains that colonize each parotid lymph node is similar between animals. These data provide evidence for a strong restriction of strains between the conjunctival mucosa and the parotid lymph node. Specifically, we observe a loss of at least two orders of magnitude in strain diversity between the site of infection and the parotid lymph nodes. This population restriction is corroborated by marked decreases in alpha and Shannon (H) diversity in the lymph nodes relative to the inoculum (**Fig 2D-E**).

### Strain abundance distribution is skewed in host tissue

The highly diverse *B. abortus* strain population in the inoculum has an even strain (i.e. barcode) abundance distribution (**Fig S6**) and a Shannon’s evenness index near 1 (**Fig 2F**). In contrast, the distribution of strains in the *B. abortus* populations isolated from bovine lymph nodes is highly skewed, whereby small numbers of strains were highly abundant, counted on the order of 10^6^ times, and the majority of codes in these tissues were rare (**Fig S6**). This skew is reflected in low Shannon evenness values for host tissues (**Fig 2F**). The most abundant strain in each lymph node accounted for 12-99% of total sequence reads (**Fig 2F**, **Dataset S1**), and in most lymph node samples 80% of strains were counted only once (**Fig 2G**). In comparison, the most abundant strain in the inoculum accounted for only 0.047% of total reads. A positive relationship between abundance in the inoculum and the likelihood that a strain will colonize the host is expected. Indeed, strains that occupy the most extreme end of the abundance tail of the inoculum are more likely to be observed in the host.

Abundant strains in technical replicates from a subset of lymph node fragments are highly correlated (**Fig S4**). This lends confidence to our conclusion that such strains are truly abundant in the lymph node and are not an artifact of sample processing or jackpot effects during PCR barcode amplification. Variation between replicates for less abundant codes reflects the limits of sampling low abundance members of a community using this method. The profound skew in the strain abundance distribution that we observe in lymph node samples provides evidence that a very limited number of strains that reach the lymph node are hyper-abundant during the early stages of infection.

### The populations of most abundant strains in each lymph node are distinct

The observed skew in the strain abundance distribution *in vivo* raised the question of whether the most abundant *B. abortus* strains were unique to each lymph node or the same across nodes. Hyper-abundance of distinct sets of strains across animals (and lymph nodes) is expected if the governing processes are stochastic. On the other hand, if certain strains have a genetic propensity to hyper-proliferate, we expect to see a shared set of highly abundant strains between animals (and lymph nodes). To evaluate these possibilities, we extracted the set of barcodes that comprised at least 2% of the sequencing reads in any lymph node sample. We then evaluated the relative abundance of these 160 barcodes in every sample. We observe a distinct set of highly abundant strains in each lymph node (**Fig 3**). There was no overlap in hyper-abundant strains between animals and, in most cases, no overlap in hyper-abundant strains between the right and left parotid lymph nodes within an animal. Furthermore, in many cases, abundant strains were distinct even between quadrants of an individual lymph node providing evidence that strains that establish residence and proliferate in the lymph node are, to some extent, spatially restricted within this tissue. However, there are cases in which abundant strains are shared among dissected quadrants of an individual lymph node. Thus, it is likely that highly proliferating strains eventually spread through the node. Indeed, the two parotid lymph nodes exhibiting the most homogeneity in abundant strains across the four quadrants (the left parotid lymph nodes of calves 3 and 4), also had the highest bacterial loads (**Fig 1 & 3**). We conclude that at early stages of infection, *B. abortus* populations are dominated by a small, random set of strains.

**Fig 3.**
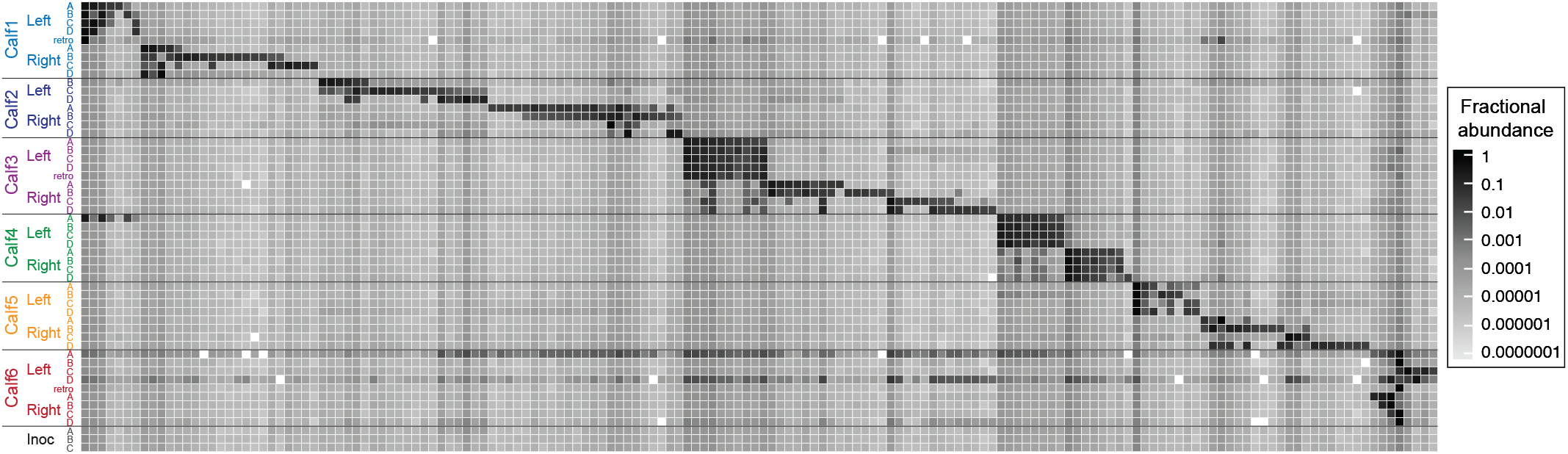
Distinct sets of hyper-abundant strains are observed in each lymph node. Heat map of the abundance of 160 barcodes that represent more than 2% of reads in at least one sample. Each row is a sample, each column represents a hyper-abundant barcode (i.e. strain), and grey scale intensity indicates fractional abundance of each barcode in each sample. Empty squares indicate barcodes that were not detected in a sample. Horizontal lines separate samples from each animal. Note that the PCR amplification of barcodes from calf 4, left parotid, quadrant A was contaminated with template from calf 1, left parotid, quadrant B. This signal is detected in the abundance data from the calf 4 Left_A sample.

### B. abortus strain populations within individual lymph nodes are spatially heterogeneous

We next sought to more comprehensively assess *B. abortus* population structure in bovine lymph tissue after infection by analyzing the full set of strains (i.e. barcodes) present in each node fragment. We utilized two different metrics for pairwise population comparisons between each sample. Jaccard dissimilarity is a metric that compares the presence or absence of population members (26). Bray-Curtis dissimilarity is a related metric that also takes into account abundance of population members (27). When considering presence/absence of barcodes in each population, we observe high levels of dissimilarity between all host tissue samples, even samples dissected from the same lymph node (**Fig 4A**). When abundance is considered, we again observe high levels of dissimilarity between populations of *B. abortus* strains from different lymph nodes in the same or different animals. Moderate similarity is detected between dissected quadrants of individual lymph nodes (**Fig 4B**). We note that the lymph nodes in which the populations in each quadrant exhibit the lowest Bray-Curtis dissimilarity (i.e. greatest similarity) tend to share more hyper-abundant strains between quadrants as well (**Fig 3 & 4B**). The largely distinct sets of strains colonizing each animal, each lymph node, and each region of a lymph node are consistent with a significant population bottleneck leading to random subsets of strains in each tissue. In addition, the spatial heterogeneity within tissues suggest that at least early in infection, most strains in lymph nodes are spatially restricted and do not spread evenly throughout the lymph node. This is consistent with granuloma formation in the lymph nodes, which is a feature typical of host tissue infected with *Brucella* (28).

**Fig 4.**
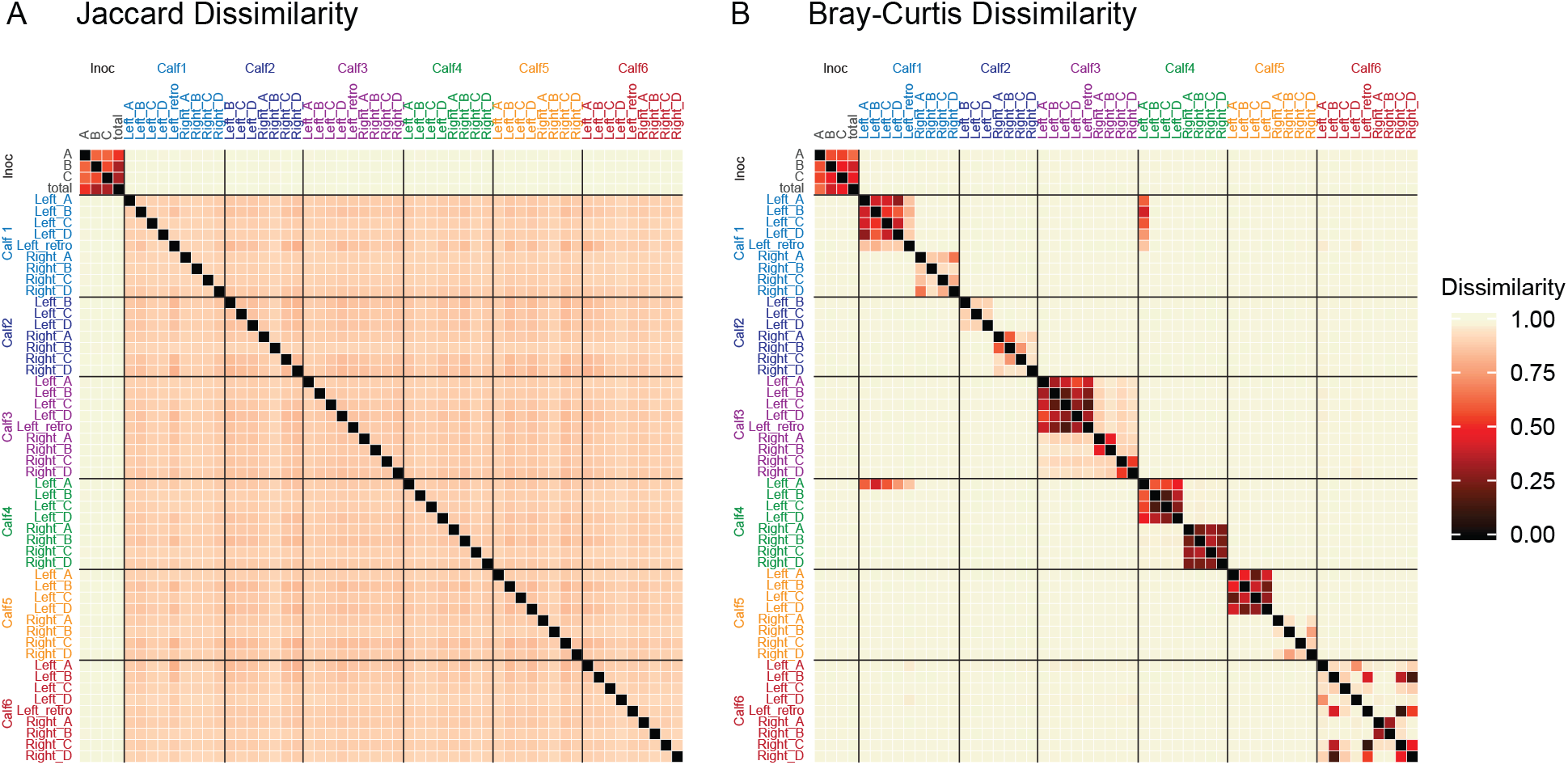
*B. abortus* strain populations are similar within a lymph node, but different between animals. Heat map of (**A**) Jaccard and (**B**) Bray-Curtis dissimilarity values for each pair of samples (1=dissimilar, 0=similar). For the inoculum (inoc), each individual replicate (A-C) and the sum of the three replicates (total) were compared. Horizontal lines separate samples from each animal. Samples are colored as in Figure 1. Again, Bray-Curtis dissimilarity scores between calf 4 Left_A and calf1 Left quadrants reflect contaminating PCR signal.

### Strain populations in retropharyngeal lymph nodes reflect the parotid lymph node in the same animal

In the three cases for which we obtained population data from retropharyngeal lymph nodes, there was a significant degree of similarity between the retropharyngeal node and its associated (draining) parotid lymph node. Specifically, the highly abundant codes in the retropharyngeal lymph node populations were shared with the highly abundant codes in the parotid lymph node from the same side of the animal (**Fig 3**), and Bray-Curtis metrics indicate population level similarity between retropharyngeal lymph nodes and the parotid lymph node from the same side of the animal (**Fig 4B**). Thus, a strain that establishes residence and proliferates in a parotid lymph node is more likely to establish residence in directly connected nodes.

### Strain-level probability of parotid lymph node colonization via the conjunctival route

Experimentally defined populations of *B. abortus* in the inoculum and in the lymph nodes of bovine hosts provide the opportunity to assess the probability of host infection at an individual strain level. There are many host immune mechanisms that function to restrict microbial entry and proliferation in the host. Some strains present in the inoculum will successfully traverse these immune obstacles and establish a colony in the host tissue, while others will be eliminated from the host. There is evidence that host entry and colonization is shaped by stochastic factors (29). However, infection is also influenced by the genetic properties of the particular strain (30). Other Tn mutagenesis studies involving *Brucella* infection in tissue culture models(31–34), or in animal models (35–38) indicate that a small fraction of mutants are attenuated. Similarly, only a small fraction of genes are required for survival of *Mycobacterium spp*. in mammalian hosts (39, 40). Thus, it is reasonable to predict that most (>90%) transposon insertions that are tolerated in axenic culture will be neutral or near neutral during infection.

If a Tn-*himar* barcode insertion at a particular chromosomal site is neutral with respect to infection, then infection can be thought of as a simple “yes-no” event that may be modeled as a binomial sampling process. In such a model, the expectation of observing a neutral barcode in the host should follow a binomial sampling distribution. Specifically, the success rate of individual strains (i.e. barcodes) in colonizing a lymph node is a function of *a*) the probability that a single cell will successfully infect and *b*) the abundance of each strain in the inoculum. In a binomial framework, the number of cells of a particular strain in the inoculum can be considered the number of infection trials for that strain.

Following this model, we empirically determined the probability that any single cell will colonize by binning the strains in the inoculum by abundance and surveying the fraction of strains in each bin that are not detected in each parotid lymph node (i.e. that are lost during infection). As expected, more abundant strains in the inoculum are more likely to be detected in a lymph node. In addition, all lymph nodes across all animals showed strikingly similar rates of strain loss in each abundance bin, demonstrating that this frequency-dependent pattern of infection is general. Furthermore, the relationship between relative abundance of strains in the inoculum and the likelihood of observing those strains in lymph nodes follows a binomial probability (**Fig 5**). We compared the experimentally determined relationship between strain abundance in the inoculum and detection frequency in the lymph nodes to theoretical expectations of binomial sampling with different probability of success. Based on this analysis, we estimate the probability that any individual cell in the inoculum will successfully colonize a parotid lymph node via the ocular conjunctiva to be 0.0001. Furthermore, fitting the experimentally observed patterns of strain loss versus colonization to a log-logistic dose response model (41) yields in an ID_50_ corresponding to fractional abundance of 10^-4.4^, or approximately 3500 CFU (**Fig S7**).

**Fig 5.**
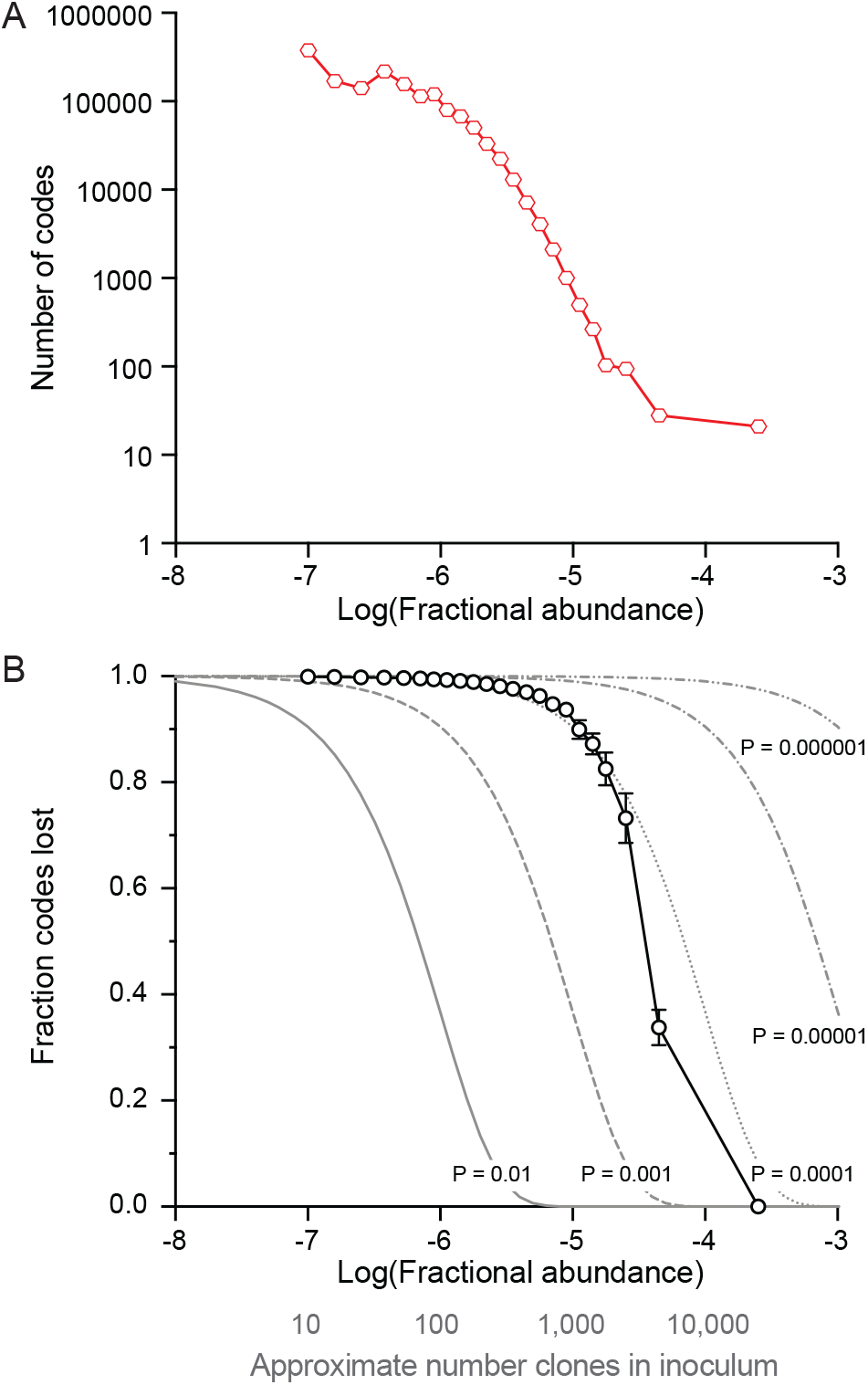
Lymph node colonization by individual *B. abortus* strains is a function of abundance in the inoculum and follows a binomial probability. **A.** Barcodes (i.e. unique strains) were binned by fractional abundance in the inoculum. The number of barcodes in each bin is plotted where abundance corresponds to the center of the bin. **B.** Fraction of strains not detected in parotid lymph node samples (i.e. lost during infection) are plotted as a function of barcode abundance in the inoculum. The presence or absence of each barcode was recorded for each parotid lymph node. Mean (± SD) fraction of codes not detected in each bin is plotted in black (n=12 parotid lymph nodes). Grey lines represent expected fractions of codes not detected assuming a binomial sampling at different probabilities (P). To simulate a binomial sampling, the fractional strain abundance in the inoculum was converted to an approximate number of strains in the inoculum (fractional abundance * 10^8^ CFU in inoculum; grey axis label) and used as the number of trials.

## Discussion

Transposon sequencing (Tn-seq) studies are most often aimed at high-throughput assessment of gene function in a particular environment (42). We constructed a barcoded pool of *B. abortus* Tn-*himar* mutants and used this population to quantify bovine colonization through a natural route of infection at the individual strain level. This work demonstrates the feasibility of applying Tn-seq approaches to study how infection shapes *B. abortus* populations in a large animal (bovine) system.

The barcoded Tn mutant collection used in this study is highly diverse (≈10^6^ unique strains; **Fig 2**, **Fig S5**). We reproducibly recovered strains from this pool from two lymphatic sites and we were able to reliably amplify *B. abortus* strain barcodes from this host tissue. The extreme stochastic loss of strain diversity between the conjunctival infection site and lymph nodes, and the corresponding skew in *B. abortus* strain abundances *in vivo* precluded a global analysis of the function(s) of individual genes during infection. Nonetheless, we were able to leverage the barcoded nature of our Tn-*himar* pool to quantify population-level features of *B. abortus* infection biology, shedding new light on an infection bottleneck and the spatial distribution of *B. abortus* clones in colonized tissue. Tn-seq studies aimed at defining the roles of individual *B. abortus* genes in bovine infection will require a less diverse infecting pool and are in development. Given the observed bottleneck and the number of CFU in the infecting inoculum, we estimate that a mutant pool containing 1000-3000 strains, each represented by 10^4^-10^5^ clones, would be suitable to assess individual gene function in the context of this infection model.

### Characterizing a bottleneck in a natural route of infection

We administered 10^8^ *B. abortus* Tn-*himar* mutants to six calves via the conjunctival route. This is 10-100 times lower than typical *B. abortus* immunization doses administered to cattle (43–45), but similar to challenge strain doses in vaccine or experimental infection studies (1, 5, 44). Thus, the number of strains delivered to calves in our study is relevant when considering the population biology of *B. abortus* in vaccination or experimental infection contexts. In a field setting, it is reasonable to expect that cattle contacting *Brucella* via abortions in the field can be exposed to 10^8^ cells (or more) given that aborted fetal tissue can contain 10^13^ CFU/gram (46).

Our dose is significantly higher than the reported *B. abortus* infectious dose of approximately 500 CFU in mice (47, 48) and guinea pigs (49), when administered via aerosol. The magnitude of strain loss that we observe between the conjunctiva and the site of initial lymph node colonization (**Fig 2**) suggests that the number of *B. abortus* cells required to ensure reliable infection from the ocular conjunctiva is much larger than 500 CFU. Indeed, controlled cattle vaccination studies (50, 51) have shown that only 78% of unvaccinated animals are infected at a dose of 3.7 x 10^5^ CFU via the conjunctival route, while 89% are infected at a dose of 7.4 x 10^5^ CFU. Doses of 1.5 x 10^7^ to 1 x 10^8^ CFU typically result in 100% infection in experimental conjunctival infections (51). From these data, an infection “bottleneck” along the conjunctival route is inferred.

Several groups have presented thorough discussions of the nature of infection bottlenecks from both experimental and theoretical perspectives (12, 52, 53). These reviews and others highlight ways in which host restriction mechanisms shape bottlenecks which, in turn, influence the population biology and evolutionary dynamics of microbial pathogens (54). Given the difference in *B. abortus* population size between the inoculum (10^8^) and the lymph node (10^5^-10^6^), a bottleneck in this path is evident. However, one cannot accurately gauge the magnitude of a bottleneck from cell count alone because population size is a function of both the bottleneck itself and subsequent microbial growth that occurs between the time of colonization and the time of sampling. By uniquely tagging individual strains in the population and leveraging the distribution of strain frequencies in our tagged pool, we were able assess the magnitude of the *B. abortus* population bottleneck via the conjunctival route of infection.

At an inoculum of 10^8^ CFU, each strain in the initial pool (diversity of ≈10^6^) was represented on average 100 times in the infecting dose. If we assume that each barcode in a lymph node reflects a single colonization event, our data indicate each node was colonized by an average of 7 x 10^3^ individuals from a pool of 10^8^ CFU in the inoculum (**Fig 2**, **Dataset S1**). Thus only 1 in 14,000 strains present in the inoculum are detected *in vivo* 7-8 days post infection; this could be viewed as an infection bottleneck of 1:14,000. This rough estimate likely undercounts colonization events because highly abundant strains in the inoculum are expected to successfully colonize more than once. As such, we extended our analysis of the probability of host colonization taking advantage of the fact that the abundance of strains in the inoculum is not completely uniform. This permitted us to assess the probability of colonization of the parotid lymph nodes as a function of strain abundance in the infecting pool. Specifically, we considered colonization at the individual strain level as a binary event, i.e. whether a strain barcode was present or absent in a node. Plotting the probability of observing a particular barcode - in any node, across all 6 animals - against the fractional abundance of barcodes in the infecting pool yielded a binomial-like probability curve, which is well modeled by a single cell colonization probability of 0.0001 across the vast majority of strain abundances. From this analysis, we conclude that the infection bottleneck at a dose of 10^8^ CFU is 1:10,000.

Multiple models can explain the bottleneck we observe. At one extreme, it could be the case that 99.99% of strains in the *Tn-himar* pool contain insertions in genes required to colonize the host. These strains would therefore be lost by the time we isolated *B. abortus* from host tissue 7-8 days after infection. Certainly, our pool of Tn-*himar* strains is expected to contain mutants that have reduced fitness in the bovine host environment. However, previous *in vitro* and *in vivo* studies with less complex *Brucella* spp. mutant pools (31–38) are not consistent with a model in which such a large fraction of mutants are attenuated in infection. In a second model, the host exerts selective pressure on all members of the infecting population irrespective of strain genotype, and only 1:10,000 cells randomly colonizes. In reality, aspects of both models influence the infection bottleneck we observe. Indiscriminate strain killing by host defenses is the dominant mechanism that accounts for loss of strain diversity in this experiment.

### *Brucella* Tn-seq studies of natural host via natural routes

This study demonstrates that barcoded Tn-seq (Bar-seq) studies provide a means to quantify changes in the population structure of *B. abortus* in a bovine host, infected via a natural route. Given the success of the approach outlined here, one can envision using similar approaches for studying *B. abortus* transmission in suitably controlled field settings. We have also recently constructed a *Brucella ovis* bar-seq mutant library (55), which may be similarly useful for assessing the population biology of this ovine pathogen during infection/transmission.

## Materials & Methods

### Bacterial strains and Select Agent considerations

All experiments using live *B. abortus* 2308 were performed in biosafety level 3 facilities according to select agent regulations at the USDA National Animal Disease Center. *B. abortus* Tn-himar strains were cultivated on Tryptic Soy Agar supplemented with 5% serum and 50 ug/ml kanamycin (TSA-SK) at 37°C with 5% CO_2_.

### Preparation of B. abortus transposon library

The *B. abortus* Tn-Himar library was previously generated and mapped (21) using a barcoded transposon mutagenesis strategy developed by Wetmore and colleagues (22). In this study, a 1 ml aliquot of the mutant library containing approximately 2.3 × 10^9^ CFU was thawed and spread on a 150 mm TSA-SK plate. After 2 days of growth at 37°C in the presence of 5% CO_2_, cells were scraped from the plate, resuspended in phosphate buffered saline (PBS), and diluted to a concentration of 2 × 10^9^ CFU/ml in PBS. This mixture of strains served as the inoculum for all 6 animals. Three 500 μl aliquots of this mixture were saved as representatives of the inoculum.

### Animal care and welfare

Holstein calves between 5 and 9 months of age (**Table 1**) housed on the grounds of the National Animal Disease Center in Ames, Iowa, were transferred to an agriculture biosafety level 3 (Ag-BSL-3) containment facility, where they were housed for the duration of the study. All animal care was performed under the guidance of the associated IACUC protocols at the National Animal Disease Center.

### Infection of cattle with B. abortus transposon library

All animal work was conducted in the BSL-3 Agricultural Suite at the USDA National Animal Disease Center, Agricultural Research Service according to USDA approved protocols. Animals were acclimated to the biosafety containment barn for one week prior to infection. 50 μl of the *B. abortus* mutant library prepared as described above was inoculated into the conjunctiva of each eye of each calf. Eight or nine days postinfection, the calves were humanely euthanized and the parotid and retropharyngeal lymph nodes were collected. The lymph nodes were immediately placed at 4°C. Parotid lymph nodes were processed the day of collection. Retropharyngeal lymph nodes were processed the following day.

### Harvesting B. abortus transposon strains from lymph nodes

Parotid lymph nodes were divided into four equal sections. Each section was weighed and then homogenized in 2 ml PBS in gentleMACS M tubes and a gentleMACS Octo Dissociator (Miltenyi Biotec) using the RNA_1 setting. Homogenates were gently mixed with detergent (0.1% Triton X-100) and incubated at room temperature for approximately 10 minutes to lyse the eukaryotic cell membranes and allow tissue debris to settle. 200 μl of the homogenate were serially diluted to enumerate CFU on TSA-SK plates. One mL of each homogenate was evenly spread on a 150 mm TSA-SK plate to recover *B. abortus* strains. Plates were incubated at 37°C in the presence of 5% CO_2_. After 72 hours, the bacterial lawn was scraped from the plate, resuspended in PBS, and cell pellets were stored at −20 °C.

Due to limitations in sample processing throughput and the expectation that retropharyngeal lymph nodes may not be colonized at the time of harvest (1), only the left retropharyngeal lymph node from each animal was processed. These were divided into two sections and homogenized as above. One ml of each homogenate was spread on a 150 mm TSA-SK plate to recover bacteria and assess colonization. After 72 hours of incubation, zero to a few *Brucella*-like colonies were observed from three animals; thousands to a lawn of *Brucella*-like colonies were observed from three animals. Bacteria were scraped from the later three samples and processed with the parotid lymph node samples.

### *B. abortus* genomic DNA prep

Genomic DNA was recovered from stored cell pellets using the Ultradeep Microbiome Prep Kit (Molzym), omitting the initial steps for removal of host DNA. Briefly, ≈20 μl in volume of each bacterial pellet (derived from lymph node scrapings) was resuspended in kit Buffer RL. Cell pellets stored from the inoculum were also processed in parallel. Bacterial cells were lysed with addition of kit BugLysis buffer at 37 °C, lysate was treated with proteinase K at 56 °C, and genomic DNA was purified using kit spin columns following a guanidine isothiocyanate treatment step. An additional ethanol precipitation was conducted on the eluates, and final pellets were each dissolved in 100 μl of ultrapure water. DNA samples were stored at −20 °C or colder prior to PCR analysis.

### PCR barcode amplification and sequencing

Transposon barcodes were amplified as described by Wetmore *et al* (22). Briefly, 20 μl PCR reactions contained 0.2-1.0 ug of genomic DNA, 0.5 μM of each primer (Barseq_P1 as the universal forward primer and an indexed Barseq_P2 primer unique to each sample (22)), and Q5 hotstart polymerase with GC enhancer (New England Biolabs). Reactions were cycled as follows: 98°C for 4 min, 25 cycles of 98°C, 55°C, and 72°C for 30 second at each temperature, then 72°C for 5 min. Amplification was confirmed by resolving 2 μl of each reaction in a 2% agarose gel. Reactions were pooled (5 μl each) and the pooled products were cleaned using GeneJet PCR purification kit (ThermoFisher). Amplified barcodes were sequenced using an Illumina HiSeq 4000 with Illumina TruSeq primers. Raw barcode amplicon sequence data have been deposited in the NCBI sequence read archive (SRA) under BioSample accession SAMN10145474; BioProject accession PRJNA604627.

### Barcode counting; a filtering approach to minimize spurious inflation of barcode diversity

Our analysis workflow is schematized in **Fig S2** and described as follows. The sequencing reads were processed with the custom perl script, MultiCodes.pl (22) to generate lists of the number of times each barcode was observed in each sample. To assess the fitness consequences of gene disruptions by barcoded transposons, the analysis pipeline developed by Wetmore et al (22) analyzes only codes that are mapped with high confidence to positions in the genome, and the pipeline to map barcodes to specific positions in the genome is stringent (22). An advantage of this approach is that it effectively minimizes the effect of sequencing errors, as codes in experimental samples that are not identical to a mapped code are ignored. However, in the analysis presented herein, we aimed to use barcodes to assess the diversity and structure of the entire population and the un-mapped codes constitute a sizeable fraction of the population (**Fig S5**). Illumina sequencing errors occur at a frequency of 10^-3^ per base (23–25), thus given a 20-bp barcode, one expects approximately 1-2% of barcode sequence reads to contain an error. Such errors inflate the apparent diversity of barcodes in a sampled population. In an effort to more accurately define *B. abortus* population distributions in host tissue, we developed the following approach to minimize the influence of sequencing errors on our analysis.

There are multiple approaches to reduce error in sequence-based population measurements (56). In our case, the *B. abortus* Tn mutant library is highly diverse and contains many true rare codes (21). Thus approaches based on removing codes under a threshold of coverage were inappropriate. We therefore applied two selective approaches to reduce inflation of barcode diversity due to sequence error. First, we systematically removed codes that were ‘off-by-one’ from any code that was counted ≥ 100 times. Based on error rates, we expect that codes sequenced more than 100 times are likely to be sequenced erroneously at least once. The most abundant codes in our host datasets were counted on the order of 10^6^ times in select lymph node samples (**Dataset S1**). For these extremely abundant codes, we detected all 60 possible codes arising from single site errors at an observed rate of 0.5-1 x 10^-3^ per site, consistent with expectations (see **Fig S1** for examples). When summed, the reads corresponding to these ‘off-by-one’ codes total 1-2% of the reads of the abundant parent code. Applying this filtering approach resulted in the removal of a small fraction (1.4 ± 0.3 %) of total sequencing reads, and 24 ± 4 % of codes from each lymph node sample. As a second step, we removed all codes from our lymph node samples that were not detected in at least one of the inoculum replicates. Together, these two filtering steps removed 80.3 ± 9.7 % of codes, which represented only 1.7 ± 0.3 % of the total reads from the lymph node samples (**Fig S3** and **Dataset S1**). Importantly, the total number of reads filtered using this approach is consistent with the expected number of erroneous reads based on known Illumina error rates. After filtering, new lists of the number of times each barcode was seen in each sample were generated. Merged lists of the barcodes derived from totaling the reads of the samples from each lymph node were also generated.

### Population analysis of barcodes

To generate the proportional Euler graphs to visualize overlap in barcode sets we used the R package eulerr (57). The R package vegan (58) was used for population diversity metrics including Fisher alpha diversity, Shannon diversity (H) and evenness (J). The vegan function rarefy was used to assess diversity in a randomly selected set of 10^6^ sequence reads. Dissimilarity between samples was calculated with the vegan function vegdist. Analysis of the hyperabundant barcodes involved first identifying barcodes that comprised at least 2% of the reads in any sample. 160 barcodes met this threshold. We then extracted the fractional abundance of this set of 160 codes in all samples.

### Probability

To assess abundance of each code in the inoculum, the counts for all three inoculum replicates were summed. The total counts for each barcode were normalized by the total number of reads in all inoculum replicates. These fractional abundance values were then log10 transformed. Barcodes were then binned by log10(fractional abundance). Most bins were 0.1 log unit in width. However, the bins for the most abundant codes were wider due to the scarcity of highly abundant codes. Next we assessed the fraction of codes in each bin that were or were not detected in the summary list of barcodes for each parotid lymph node. Then for each bin, we averaged the fraction not detected in all 12 lymph nodes.

To generate the probability curves, we used the binomial distribution function [*p*(*x*) = choose(*n,x*) *P^x^*(1-*P*)^(*n-x*)^] to calculate the probability (*p*) of colonization occurring exactly zero times (*x* = 0) with different probabilities (*P*) of colonization over a range of number of trials (*n*) using the pbinom function in R. Here the number of trials corresponds to abundance in the inoculum. Specifically, *n* was calculated by multiplying fractional abundance in the inoculum by the size of the inoculum (10^8^).

To estimate the infectious dose at which 50% of lymph nodes would be colonized (ID_50_), the observed mean fraction of strains not detected in lymph nodes was fit to a log-logistic model with three parameters using the drm function in the drc package in R (41). In this model, *f*(*x*) = *d*/(1+exp(*b*(log(*x*)-log(*e*)))), where *d* is the upper limit, *b* corresponds to the steepness of the curve, and *e* indicates the dose (i.e. fractional abundance) corresponding to a 0.5 probability of success in colonization.

## Supporting information

Supplementary_Datatable

## Acknowledgements

We wish to acknowledge the efforts of the project technicians, especially Darl Pringle, the animal care and veterinary staff, and the biosafety staff at the National Animal Disease Center. We thank David Hershey, other members of the Crosson lab, and Andrew Olive for helpful discussions. This work was funded by in part by NIH R01AI107159 and R35GM131762.

