## Supplementary_Figures for "Quantification of *Brucella abortus* population structure in a natural host"

**This PDF file includes:**

Figures S1 to S7

**Other supplementary materials for this manuscript include the following:**

Datasets S1

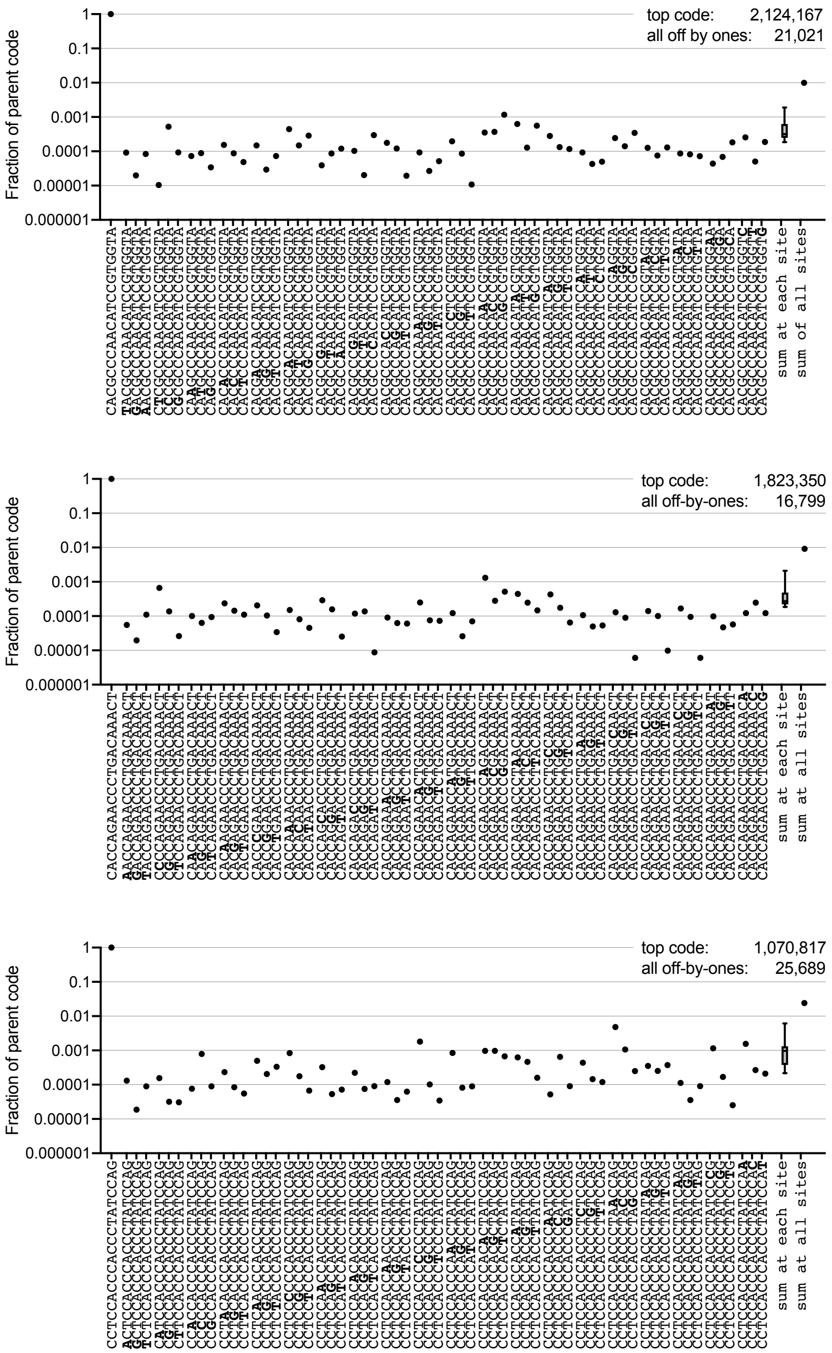


**Fig S1. Abundant barcodes reveal sequencing error rate.**

Barcodes with one difference from the most abundant 20-bp barcode in three example samples: 1_Left_A, 4_Right_A and 5_Right_C (top to bottom). The abundant code is on the left, followed by each of the 60 possible ‘off-by-one’ codes with the changed base in bold. The fractional abundance of each code relative to the parent code is plotted. Box-plot represents the range and mean of the sum of three ‘off-by-one’ codes at each of the 20 sites. Finally, the sum of all 60 ‘off-by-one’ codes is plotted. The number of reads corresponding to the abundant parent code and the 60 ‘off-by-one’ codes is indicated at the top right of each plot.

1. **Initial processing - all samples**
Extract and count barcode sequences from raw fastq files using `multicodes.pl`

2. **Filtering - Inoculum samples**
In each sample:
• ID abundant barcodes (≥100 counts)
• Remove barcodes that are "off-by-one" from abundant barcodes
→ Barcode list, with counts, for each inoc sample

• Sum filtered reads for three inoculum replicates
→ Total inoculum barcode set with total counts

3. **Filtering - Lymph node (LN) samples**
In each sample:
• ID abundant barcodes (≥100 counts)
• Remove barcodes that are "off-by-one" from abundant barcodes
• Remove barcodes absent in total inoculum list
→ Barcode list, with counts, for each LN fragment

• For each LN, sum filtered reads from each quadrant
→ Barcode list with total counts for whole LNs

4. **Analysis - Population diversity metrics**
• Calculate Fisher's alpha diversity, Shannon diversity and Shannon evenness from each sample using `vegan` R package.
→ Figure 2

• Calculate dissimilarity between all pairs of samples using `vegdist` function in `vegan` R package.
→ Figure 4

5. **Analysis - Hyperabundant codes**
• ID hyperabundant barcodes (>2% of reads in any sample)
• Determine fractional abundance of these barcodes in all samples
→ Figure 3

6. **Analysis - Probability of colonization**
• Bin barcodes in total inoculum list by log(fractional abundance)
• In each bin of inoculum barcodes, score presence / absence in each whole LN list.
• For each bin, average fraction absent in each of the 12 LNs.

• Calculate expected fraction absent from LNs assuming binomial sampling with different probabilities of success using `pbinom` function in R.
→ Figure 5

• Fit mean to log-logistic model with 3 parameters to estimate $ID_{50}$.using '`drm`' function in the `drc` R package.

**Fig S2. Sequence data processing procedure outlining filtering and analysis steps.** See Materials & Methods in main text for more details.

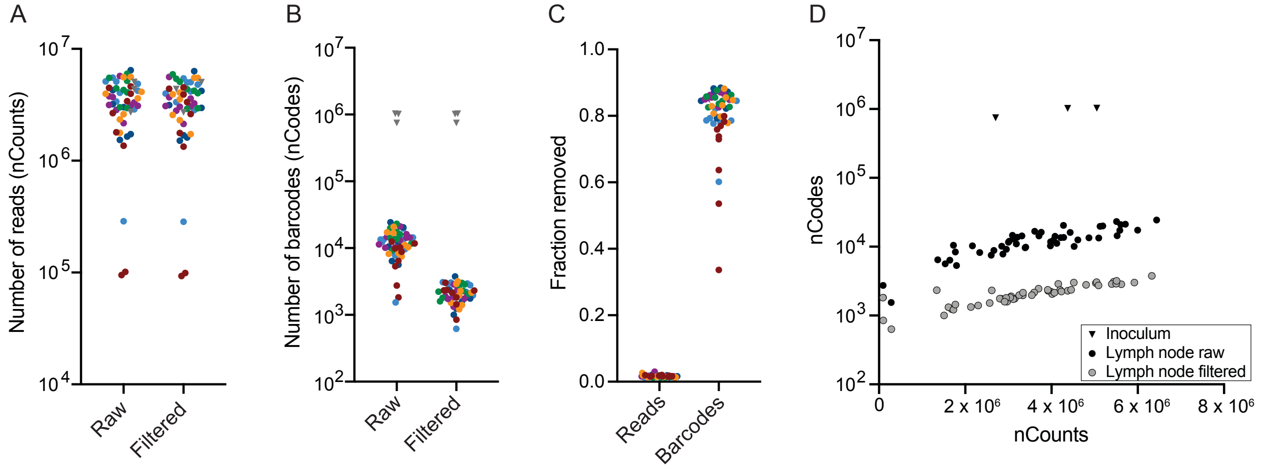


**Fig S3. Filtering to remove barcodes that are "off-by-one" from abundant (≥100 counts) codes and absent in inoculum.**

**A.** Total number of barcode sequence reads and **B** unique barcodes in each sample before and after filtering. **C.** Fraction of reads and codes removed from each sample. Samples are color coded by animal (see main text). Inoculum samples are grey inverted triangles. **D.** Scatter plot of barcodes (nCodes) per read (nCounts) before (black circles) and after (grey circles) filtering for each lymph node sample. The change in inoculum replicates (inverted triangles) upon filtering is undetectable at this scale. See **Dataset S1** for underlying data from each sample.

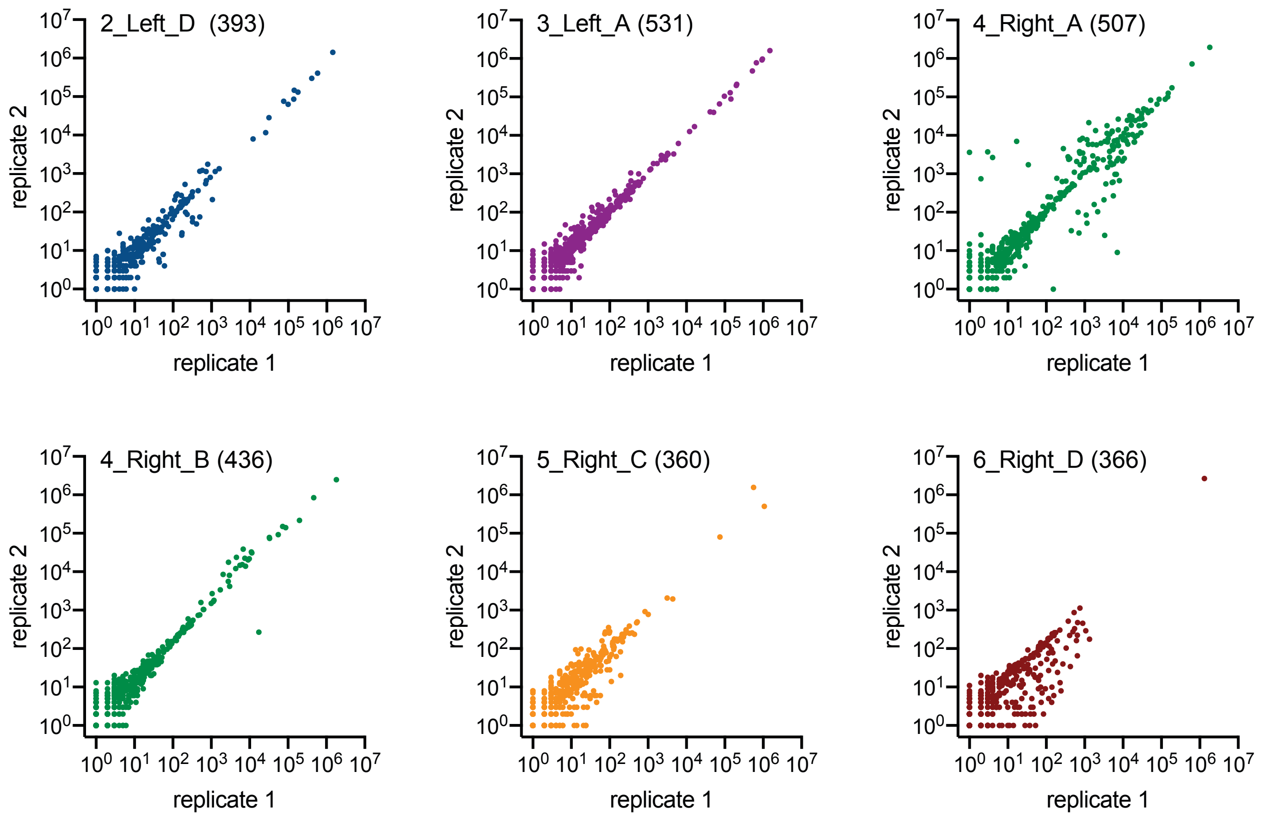


**Fig S4. Abundant codes are highly correlated between replicates from the same lymph node samples.**
For a subset of samples (indicated in the top left of each graph), we extracted two genomic DNA samples from the harvested cells and independently amplified barcodes from each replicate. The number of reads corresponding to each barcode in the two replicates is plotted where each point corresponds to a barcode. The number of shared codes is indicated in parentheses. Replicate 1 was arbitrarily selected for all subsequent analysis in this work.

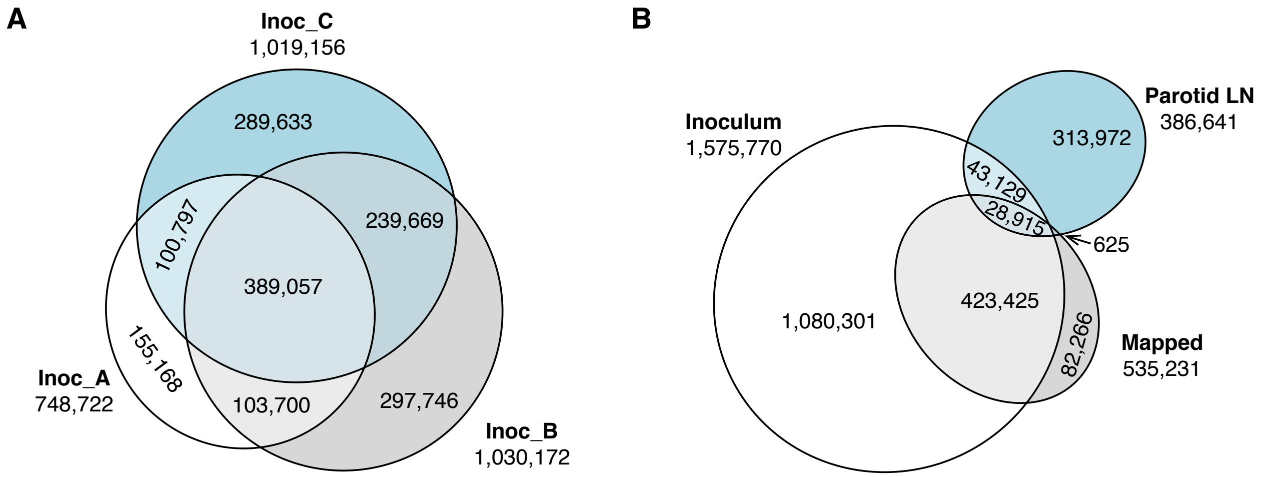


**Fig S5. *B. abortus* Tn-*himar* strain identity overlap.**
Each Tn-*himar B. abortus* strain in the pool used for animal infection (i.e. the inoculum) contains a unique 20 base pair barcode that can be used for strain identification. **A.** Proportional Euler diagram of the unique barcodes detected in each of the three replicate inoculum samples (A, B & C). The total number of codes detected in each replicate are indicated outside the circles. **B.** Proportional Euler diagram of the set of barcodes detected in the inoculum samples, the set of barcodes detected in parotid lymph node samples, and the set of barcodes previously mapped (1) with high confidence to positions in the *B. abortus* genome. The number of codes in each set are indicated outside the circles.

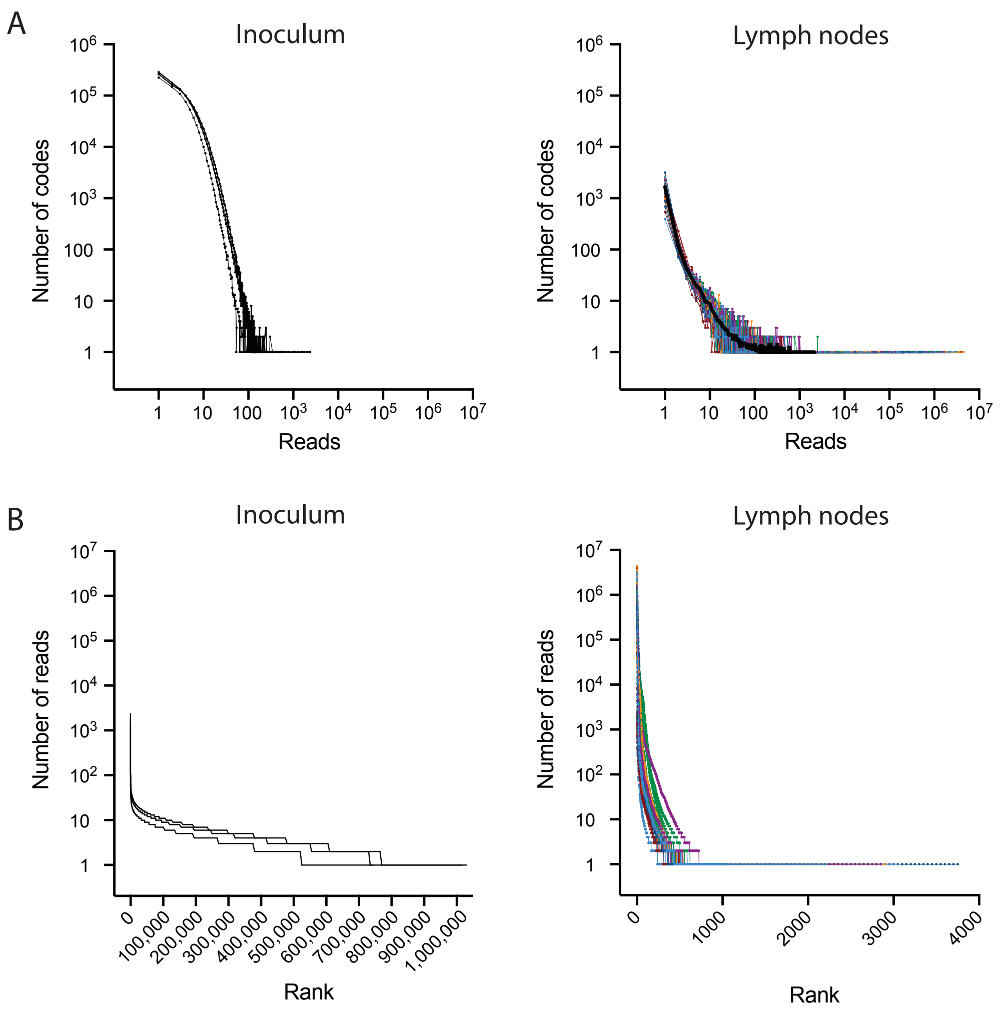


**Fig S6. *B. abortus* strain (i.e. barcode) abundance distributions in the inoculum and in bovine lymph nodes after conjunctival infection.**

**A.** Frequency of barcode abundances in each of the three inoculum replicates (left) and bovine lymph node samples (right). **B.** Ranked barcode abundance showing the number of Illumina sequencing reads corresponding to each barcode. Barcodes are ranked from most to least abundant in each inoculum sample (left) and lymph node sample (right). In all panels, each line represents the distribution in a single inoculum replicate or in a single lymph node quadrant. The lymph node samples are color coded by animal as in the main text. In (A), black line in lymph node plot represents an average of all lymph node samples.

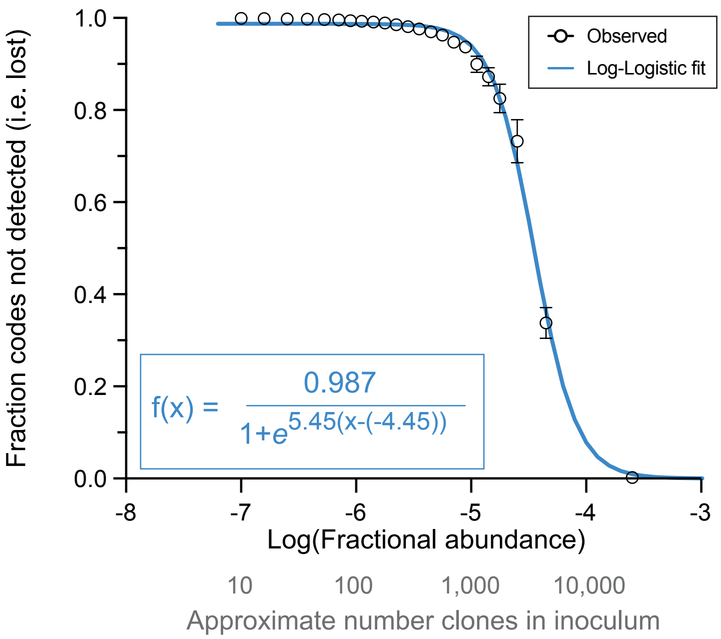


**Fig S7. Estimating infectious dose from the observed pattern of abundance-dependent loss vs. colonization.**
Observed fraction of strains not detected in parotid lymph node samples (i.e. lost during infection) as a function of abundance in the inoculum (from Figure 5 in the main text) was fit to a log-logistic model with 3 parameters: f($x$) = $d$/(1+exp($b$(log($x$)-log($e$)))). Here $x$ and $e$ are in log space, thus f($x$) = $d$/(1+exp($b$(x-$e$))). In this model, $b$ relates to the steepness of the curve, $d$ represents the upper asymptote and $e$ denotes the effective dose ($ED_{50}$), or in this case, the infectious dose ($ID_{50}$) (2). The fit parameters are $b$ = 5.45 ± 0.28 p=1.96e$^{-14}$; $d$ = 0.987± 0.004 p < 2.2e$^{-16}$; $e$ = -4.45 ± 0.01 p < 2.2e$^{-16}$. Thus, the $ID_{50}$ corresponds to a fractional abundance of $10^{-4.45}$ = 3.5 x $10^{-5}$ or approximately 3500 clones in our inoculating dose.

~~~~~~~~~~~~~~~~~~~~~~~~~~

**Dataset S1 (see separate excel file): Basic sample parameters including number of reads and number of barcodes counted before and after filtering.**
See tab 1 for key to columns. See tab 2 for data.

~~~~~~~~~~~~~~~~~~~~~~~~~~
